# Flu Mutation Explorer: an Interactive Platform for Mapping Host Adaptation Mutations in Influenza A Viruses

**DOI:** 10.64898/2026.07.22.740012

**Authors:** Laura Mojsiejczuk, Derek Wright, Robert J Gifford, Thomas P Peacock, David L. Robertson, Joseph Hughes, Daniel Goldhill, Edward Hutchinson

## Abstract

A rapid expansion of influenza A virus (IAV) genome sequencing has transformed global surveillance but has also created major challenges for interpreting the biological significance of viral mutations, particularly amino acid replacements associated with host adaptation. Resources have been created to support mutation annotation and phylogenetic analysis, but there is a need for a tool that integrates experimentally derived phenotypic evidence with evolutionary context in a framework suitable for users without prior training in bioinformatics. Here, we present the Flu Mutation Explorer, an interactive web application that combines large-scale influenza phylogenies with a manually curated database of reported mammalian adaptation mutations, to enable the exploration and interpretation of IAV genetic variation. The underlying database comprises over 1.5 million publicly available IAV sequences and over 1000 mutations associated with mammalian adaptation. The Flu Mutation Explorer enables users to query protein sequences, visualise amino acid distributions across viral lineages, examine host-specific conservation patterns, and identify adaptation mutation with links to supporting literature. We include case studies which demonstrate the platform’s use in assessing amino acid conservation at sites of interest and in rapidly identifying candidate mammalian adaptation mutations during the ongoing H5N1 panzootic. By integrating genomic, phylogenetic, and functional information into an intuitive interface, the Flu Mutation Explorer lowers the barriers to interpreting influenza sequences for specialists and non-specialists alike.

## Introduction

Influenza A viruses (IAVs) are major pathogens of humans and of many wild and domesticated animals [1]. IAVs have segmented RNA genomes, which evolve rapidly through the accumulation of mutations and through reassortment between strains. This evolution can be driven by selective pressures, including host immunity and the use of antivirals [2]. IAVs also have the capacity to adapt to new host species, a property which has caused numerous pandemics [3]. The genetic changes associated with altered host tropism are of key public health importance, and timely identification of these mutations is needed to support pre-pandemic risk assessments.

Decades of intensive global surveillance have generated vast quantities of IAV sequence data. This creates unprecedented opportunities to study viral evolution and to address public health concerns such as the adaptation of avian IAVs to mammalian hosts. However, the sheer volume of IAV sequences also makes these analyses challenging to perform, particularly for non-specialist users.

The ability to rapidly identify and interpret mutations is particularly important for public health surveillance and pandemic preparedness. Several national and international agencies actively track IAV mutations. The World Health Organisation’s Risk Assessment Tool (TIPRA) [4] and the Centers for Disease Control (CDC)’s Influenza Risk Assessment Tool (IRAT) [5] provide frameworks for evaluating the pandemic risk posed by emerging influenza viruses, and public health agencies such as the UK Health Security Agency (UKHSA) report risk assessments of avian influenza for human health [6]. Mutation monitoring efforts also contribute to vaccine composition discussions held biannually through the Global Influenza Surveillance and Response System (GISRS) [7]. Together, these activities highlight the importance of rapidly identifying and interpreting mutations associated with host adaptation and antigenic change.

Multiple tools exist to support the identification and interpretation of mutations in influenza virus genomes and offer valuable features such as mutation annotation, literature-based phenotypic interpretation, and reporting of temporal and geographic patterns. For example, FluSurver provides detailed functional insights from curated literature [8], FluPhenotype reports antigenic predictions, host predictions and mammalian virulence-related and drug-resistance-associated amino acid markers [9], and FluMut focuses on mutations affecting H5N1 viruses [10]. The Influenza Mutation Checker [12], which provides a command-line script for screening proteins against a user-defined mutation panel, was originally developed to identify mutations listed in the CDC H5N1 Genetic Changes Inventory. Nextclade [11] provides clade assignment, mutation calling, and sequence quality checks, while Nextstrain allows for real-time tracking of the evolution of multiple pathogens including specific lineages of IAV (e.g., H5N1 cattle outbreak, and H3N2 seasonal influenza). Despite their value for specific tasks, these tools do not effectively integrate experimentally determined phenotypic effects of mutations with their phylogenetic context. This limits their ability to capture patterns of clade-specific adaptation or of the emergence of mutations within a lineage. Hence, despite the importance of the tools developed so far, there is a need for an accessible tool for researchers and surveillance laboratories to easily explore IVAV mutations of interest within their broader evolutionary context.

To address this gap, we developed the Flu Mutation Explorer, a flexible, user-driven Shiny application that integrates IAV phylogenetic trees with systematically curated tables of mutations associated with mammalian host adaptation, to support the visualisation and interpretation of mutations in IAV proteins. By integrating genomic data with functional annotations and phylogenetic context, the Flu Mutation Explorer bridges the gap between surveillance pipelines and exploratory research tools. It is designed to be user-friendly, free to access, and customisable, providing a resource that supports the rapid investigation of adaptation-associated mutations in any influenza dataset by all users, including those lacking specialist bioinformatics training.

## Methods

### Database Assembly

To assemble the sequence database, IAV nucleotide sequences and associated metadata were retrieved from the NCBI Entrez databases using the IAV taxonomic ID (txid11320) with the E-utilities API [12]. Retrieved sequences were then compared using BLASTN [13] against a custom database made of a curated IAV reference set comprising the eight viral segments. This step was used to (i) identify the closest references for each sequence, (ii) validate the segment number, and (iii) exclude non-IAV sequences. Based on these results, sequences were organised into segment-specific datasets and processed as follows: first, Nextalign [11] was used to generate sub-alignments containing sequences that shared the same closest IAV reference identified in the BLAST step; second, a reference alignment of the IAV reference set (available in the project repository https://github.com/centre-for-virus-research/Flu-Mutation-Explorer), was used to guide gap insertion in each sub-alignment, ensuring consistency with the corresponding reference sequence; finally, sub-alignments were concatenated to obtain a full alignment for each segment. Nucleotide alignments were subsequently trimmed to the coding sequences (CDS) of the IAV proteins and translated into amino acid alignments.

### Sequence Curation

Several curation steps were introduced during database assembly: removing non-IAV genomic sequences, excluding sequences with no significant BLASTN hit, removing sequences shorter than a pre-established length threshold for each segment (minimum length for segments 1 -8: 2270, 2270, 2150, 1680, 1500, 1400, 980 and 840 nucleotides, respectively), and excluding sequences that could not be aligned to their corresponding reference sequence with Nextalign. Metadata associated with the sequences were curated and validated in parallel. This included the standardisation and validation of host, parsing of subtype and strain name fields, as well as the removal of duplicated records based on strain names.

### Sequence Clustering for visualisation

For visualisation and to avoid biases linked to extensive sequencing of seasonal strains, the sequence database was clustered based on similarity at the nucleotide sequence level using MMseqs2 with a 95% sequence identity threshold [14]. Representative sequences for each cluster and segment were then extracted from the nucleotide full alignments. Maximum-likelihood trees were inferred with IQ-TREE (v2.1.2) using the best-fit model and were midpoint-rooted for visualisation purposes [15, 16].

### Curation of Adaptation Mutations

Mutations reported to be associated with human adaptation of avian influenza viruses were identified through a combined literature mining and database curation approach. Missense mutations were first compiled from existing influenza mutation resources [8, 10, 17]. A structured literature search was then conducted using Google Scholar and Litsense, a sentence level searching tool from NCBI. Segment-specific queries (e.g., “PB2 mutation” and “adaptive mutation in PB2”) were performed iteratively for each segment, alongside broader searches including “*Influenza & “human adaptation”* and *“Influenza adaptive H5N1*.” For Litsense, sentence-level queries were adapted per segment (e.g. “This PB2 mutation is adaptive in humans” and “This PB2 mutation is adaptive in mammals”), replacing PB2 with each protein name. A ‘snowball’ approach was also taken to identify additional papers of interest from reference lists. Mutations were additionally extracted from studies performing deep mutational scanning (DMS), including variants not yet observed in natural populations. For each DMS dataset, a study-specific effect size threshold was applied to define significant mutations: for HA, an effect size >1 for increase in α2,6-linked sialic acid binding and pH stability [18]; and for PB2, an effect size >1 in A549 and >1.5 for mutational differential selection between human and avian cells [19]. Mutations associated with mammalian adaptation were retained where there was no evidence of reduced fitness in humans, and all curated mutations were compiled into a unified dataset for the web application.

### Interactive Web Interface

The web application was developed using the R Shiny framework (v1.12.1) [20] and deployed on https://flu-me.cvr.gla.ac.uk/

Pre-processing of the database outputs was performed to enable efficient queries at runtime, including amino acid lookup tables for position-based queries, and reference mapping tables for segment- and subtype-specific views.

The application interface is organised into two functional modules.

The Tree module renders interactive phylogenies using the Taxonium React component (v2.0.227) [21], allowing users to explore sequence diversity and associated metadata for each of the eight IAV segments. Users may query the database in two ways: by submitting an amino acid sequence, which triggers an automated BLASTP search for segment autodetection, followed by a segment-specific database search to highlight the closest representative on the tree; or by querying a specific amino acid position. For the haemagglutinin segment, positions are reported based on mature peptide numbering, whilst for all other segments, full-length protein numbering of the longest protein per segment is used.

The Adaptation Mutations module displays a curated catalogue of mammalian mutations reported to be associated with adaptation for each segment. Users may browse the full catalogue or submit a query amino acid sequence; in the latter case, the segment is autodetected as above and the query is aligned to the corresponding reference sequence using pairwise alignment. Identified mismatches are cross-referenced against the adaptation mutation catalogue, and a stacked bar chart shows the frequency of the observed amino acid at the corresponding consensus position across host groups in the database. It is important to note these mutations may only be adaptive in a specific sequence context and any ‘adaptation mutations’ detected in a novel sequence should be considered putative.

Amino acids colours are standardised throughout the application based on chemical properties, with those sharing chemical properties having different shades of the same colour. For segments encoding multiple proteins, both the Tree module and the Adaptation Mutations module handle only the major protein, i.e. PB1 (segment 2), PA (segment 3), M1 (segment 7), and NS1 (segment 8).

Finally, a Home tab displays a database status summary including last update, the number of sequences analysed and the number of sequences retained after curation and clustering per segment.

## Results

### Alignments and Phylogenies

To create Flu Mutation Explorer, we downloaded all available IAV nucleotide sequences from NCBI (1,594,392 sequences as of June 8th, 2026). Following the curation steps, a total of 1,179,380 sequences remained, with the largest proportion of sequences corresponding to the HA segment, reflecting its central role in influenza surveillance and sequencing efforts.

Sequence alignment was a critical and technically challenging step given the number and genetic diversity of sequences. To address this, we implemented a two-step alignment strategy. Sequences were first aligned against their closest IAV reference using Nextalign, and sub-alignments were subsequently merged using a curated master reference alignment to ensure positional consistency across the full dataset. This approach required the curation of segment-specific reference sets, in which lineage representatives were selected from comprehensive phylogenies of each segment to guarantee broad and balanced coverage of IAV genetic diversity and maximise alignment accuracy for any query sequence. Reference set alignments are publicly available in the project repository.

The number of representative sequences following clustering substantially reduced the dataset for faster phylogenetic tree building and improved visualisation, to 0.92% of total sequences for NA, 0.62% for HA, and 0.38 – 0.1% for the remaining segments (Supplementary Figure S1). Between February 16^th^ and June 8^th^ 2026, the number of representative sequences increased by 76,889. This corresponded to a moderate and consistent increase across all segments, with growth rates ranging from 1.7% (Segment 5) to 15.5% (Segment 3). Sequences encoding surface glycoproteins were added at different rates over this period, with NA growing 13.8% while HA grew only 5.1%.

### Mammalian adaptation mutations

A total of 1179 unique mutations associated with mammalian adaptation were compiled across all 8 segments, with most mammalian adaptations found in the HA and PB2 segments (Figure 1a). Mutation data were derived from 234 papers (available at https://github.com/centre-for-virus-research/Flu-Mutation-Explorer; Figure 1b). Of the 1179 missense mutations, 619 were identified from DMS studies (the most prominent studies in Figure 1b), which mainly identified sites in PB2 [19] and HA [18]. Overall, mutations mapped to 666 unique sites, representing approximately 2 – 12% of the possible sites in each segment (Figure 1c). The large number of potentially adaptive mutations in HA and PB2 was expected as avian viruses have previously acquired mutations in these segments when adapting to humans [22]. Most research has therefore focused on these segments. NA and PB1 had the fewest number of unique sites reported in the literature, when adjusted for segment length. PB1 is highly conserved and a higher percentage of sites in this segment are likely to be constrained. The lack of reported adaptation in NA may be an artefact reflecting the reduced research focus on NA compared to HA. NA has a relatively large region (sites 120-275) with no reported mammalian adaptations.

**Figure 1:**
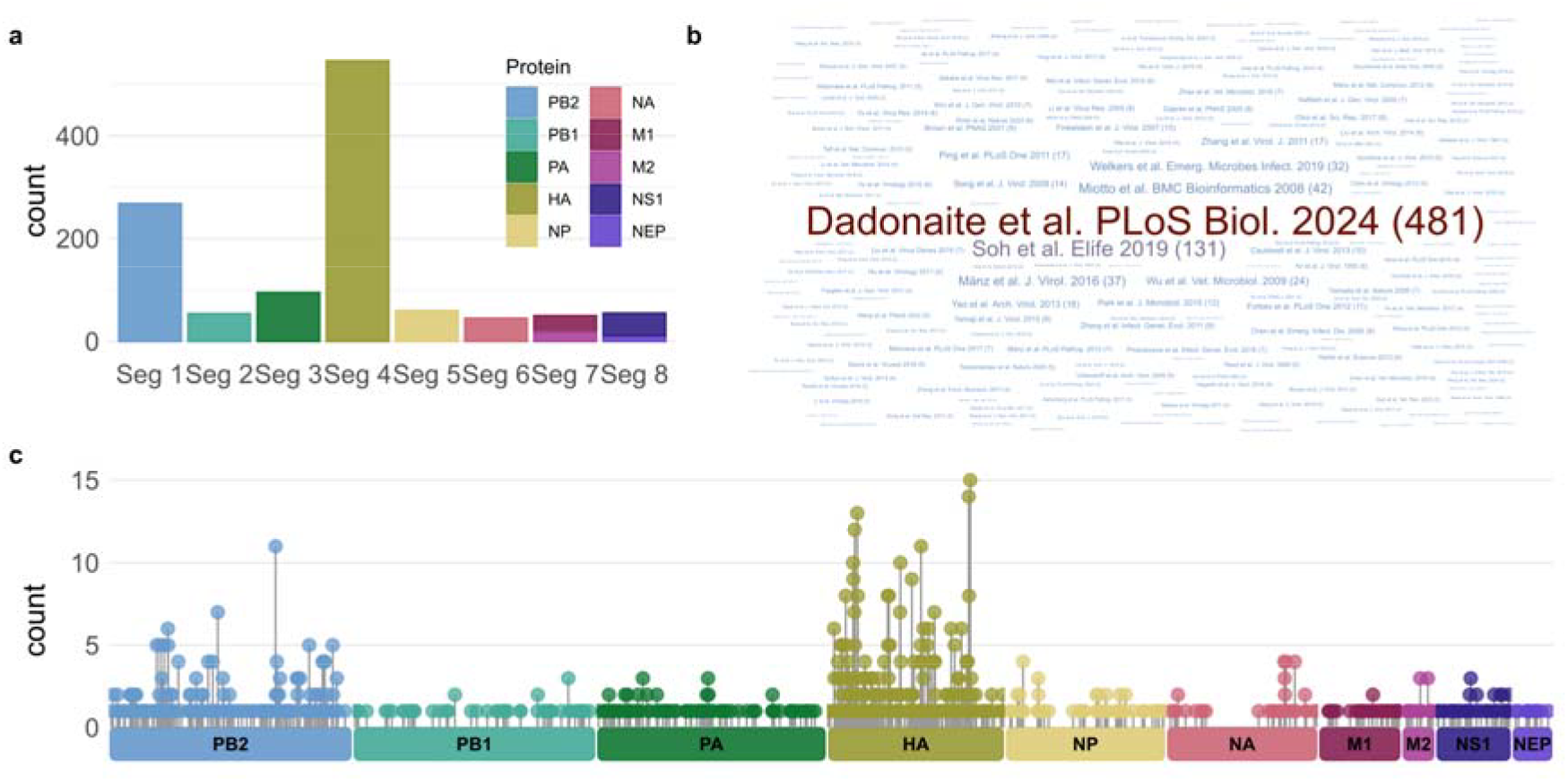
Overview of reported adaptive mutations collected from the literature. **(a)** Number of adaptive mutations per genome segment, coloured by protein. **(b)** Word cloud of publications contributing to the adaptive mutations catalogue; text size is proportional to the number of mutations reported per study and the number in parentheses indicates the number of mutations reported by each study. **(c)** Distribution of mutations along each protein sequence, with heights corresponding to the number of times a specific amino acid change in that position was reported across studies.

### Case Study: Assessing Influenza A Virus Site Conservation

The Flu Mutation Explorer provides a user-friendly tool for exploring the evolutionary context of IAV amino acid residue positions of interest. It handles otherwise onerous data curation and alignment tasks, converts between the different residue position numbering conventions used for HA and NA subtypes, and provides customisable, publication-ready figures.

For an example of the application’s basic functionality, we referred back to a previous study, in which one of us (EH) mapped the phosphorylation sites of influenza viruses using mass spectrometry [23]. Residues in proteins from the H1N1 IAV strain A/WSN/33 had been shown experimentally to be phosphorylated, and to determine if these sites were conserved a simple assessment was made of whether residues capable of phosphorylation (serine, threonine or tyrosine) were conserved at homologous positions in other IAVs. However, this analysis was limited. As the author in question did not have a bioinformatics background, the analysis was restricted to overall conservation of IAV residues and did not extend to lineage specific conservation patterns. Even that simple analysis would now be difficult to perform, as a dramatic increase in publicly available IAV sequences since the original publication (approximately ten-fold) has made it significantly harder for non-specialists to download and align IAV sequence data.

With the Flu Mutation Explorer, it is now straightforward to rapidly assess the conservation of sites of interest from the original paper (Figure 2). For each position of interest, we first annotated the IAV phylogenetic tree with residue usage at that position (Figure 2a). A significant advantage of this approach is that the Flu Mutation Explorer allows the confident identification of homologous sites in divergent IAV lineages. This is particularly valuable for highly variable proteins such as HA, for which the Flu Mutation Explorer allows seamless conversion between the numbering systems for different HA subtypes, including straightforward comparisons of Group 1 and Group 2 HA sequences (Figure 2b). We adopted the Group 1 HA residue numbering (excluding the signal peptide region) as our default reference, and maintained the alignment coordinates for Group 2. This enabled conversion between Group 1 and Group 2 numbering systems by choosing the preferred reference, and direct comparisons across groups [24].

**Figure 2:**
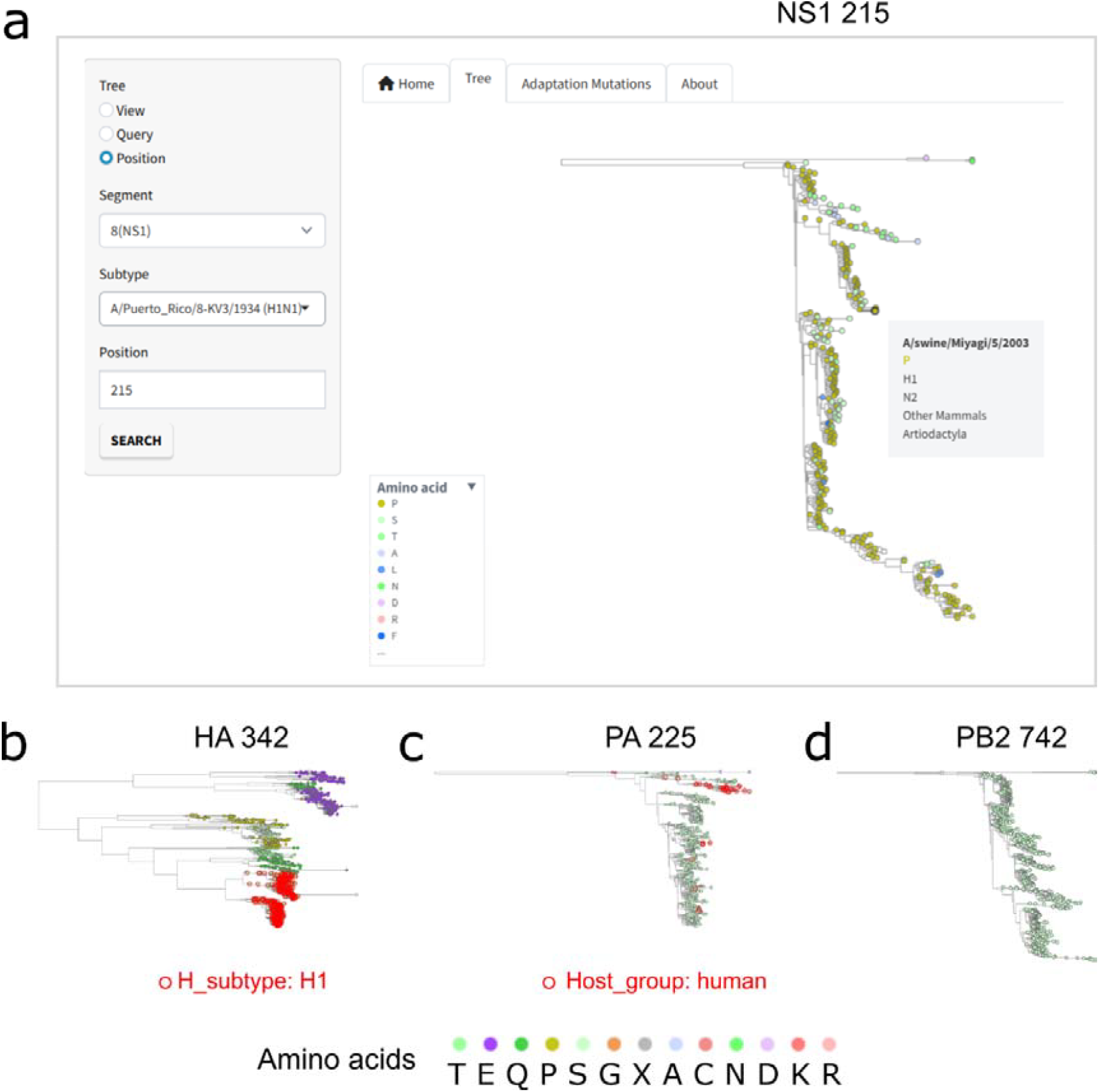
Visual inspection of residue conservation. A case study in which the Flu-Mutation Explorer was used to assess the conservation of phosphorylation sites previously identified in the IAV strain A/WSN/33 (H1N1). Phosphorylation can take place at S, T or Y resides. Figures show trees for three different IAV genome segments, with amino acid usage at the indicated positions. **(a)** A screenshot of the resource being used to assess position 215 of the NS1 protein, a residue which is not highly conserved. A grey box shows details of a highlighted strain as mouseover text. Similar analyses identified positions that are **(b)** conserved within the H1 subtype (HA 342), **(c)** conserved within human hosts (PA 225), or **(d)** absolutely conserved (PB2 742). In (b) and (c) the search terms used to highlight nodes with red circles are shown. In (b) the alignment has correctly identified homologous positions in the divergent Group 2 and Group 1 HA lineages (the major lineages in the upper and lower parts of the tree, respectively).

Using the Flu Mutation Explorer, when we observed variation at a residue, we could inspect details of strains which varied using mouse-over text (Figure 2a). Doing this helped us to identify patterns of variation that could be used to develop hypotheses about residue usage at particular sites - for example, certain residues appeared to be conserved within particular host species, while others appeared to be conserved within viral subtypes. To test these hypotheses, secondary annotations were applied to the phylogenetic trees, using rings to highlight strains with a specified trait. In this way we rapidly established detailed context for sites of interest. This allowed us to test hypotheses about the possible significance of residues based on their conservation, information which could then be used (as in our previous paper) to direct further mechanistic studies. In this case, a lack of conservation indicated that phosphorylation at a site was unlikely to be of functional importance (Figure 2b), conservation in a specific lineage indicated that if phosphorylation is important it would only be relevant in a particular genetic background (Figure 2c), and conservation across all lineages suggested that phosphorylation at this site might be functionally important for all IAVs (Figure 2d). Finally, we were able to export publication-quality vector or bitmap (SVG or PNG) files showing these analyses at the click of a button (Figure 2).

### Case Study: Identifying Mammalian Adaptations in an Ongoing Influenza Panzootic

A specific application of the Flu Mutation Explorer is to identify residues associated with the mammalian adaptation of avian influenza viruses. Here, we illustrate this for an influenza virus outbreak that was ongoing as the resource was developed.

In 2024, an outbreak of highly pathogenic avian influenza A(H5N1) was detected in dairy cattle across the United States of America, marking the first sustained transmission of this virus documented in cattle. Initial reports of viral genome sequences did not indicate mutations associated with mammalian adaptation [25]. The outbreak in cattle was part of a wider H5N1 outbreak and had complex transmission dynamics, with the virus not only spreading among cattle but also exhibiting spillover events into poultry, wild bird populations, various mammalian species, and humans. This emerging situation created a need to rapidly screen newly released viral genome sequences for signatures of mammalian adaptation.

To do this, we took the first released sequences from cattle, cats and peridomestic birds and used the Flu Mutation Explorer to check for potential mammalian adaptations. This revealed two mutations. PB2 M631L was present across all sequenced bovine isolates but had not previously been noted on an H5N1 background, and PA K497R was observed in most sequenced strains in the outbreak (Figure 3). Identifying two mammalian adaptations in these sequences, which were not present in the nearest pre-outbreak avian ancestor, strongly suggested mammalian adaptation within cows or spillovers into cats and peridomestic birds. This hypothesis was supported by additional intermediate sequences which contained M631L alone, as well as a human sequence basal to the cattle outbreak which showed different adaptive mutations in both PB2 (E627K) and PA (K142E).

**Figure 3:**
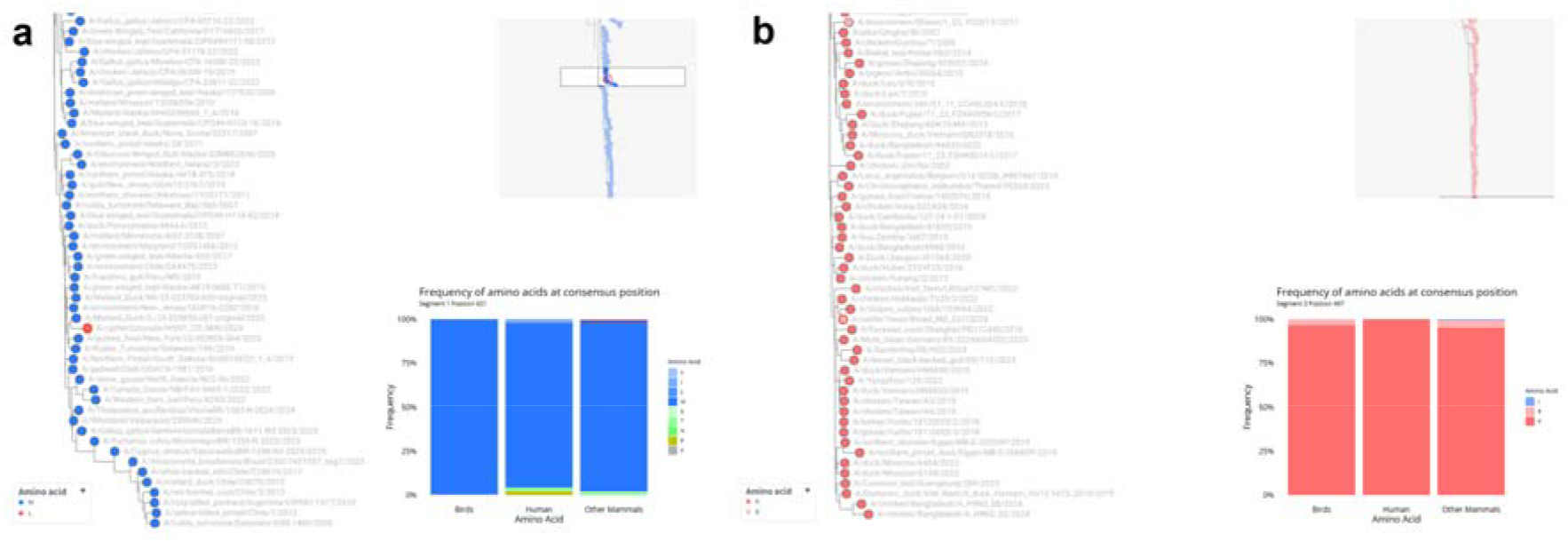
Host-specific adaptive mutations in H5N1 polymerase genes. Examples of phylogenetic trees exported directly from Flu Mutation Explorer (as PNG files) showing amino acid usage at two critical polymerase positions in H5N1 influenza viruses following cross-species transmission to cattle. Trees display amino acid residues at **(a)** PB2-631 and **(b)** PA-497, with cattle isolates (circled in red) demonstrating distinct substitutions compared to the ancestral avian lineage. Position PB2-631 shows the ubiquitous M631L substitution in all bovine isolates, while PA-497 exhibits the K497R substitution in the majority of cattle-derived sequences. Bar charts illustrate amino acid frequencies across host groups (birds, cattle, and other mammals)

We then applied the Flu Mutation Explorer to sequences that were released as the outbreak continued. We observed multiple polymerase mutations arising in sequences from cows, including PB2 E627K, D701N and D740N. However, we did not observe any human-adaptive mutations in the HA, which reduced our concerns about immediate sustained spillovers to humans. Interestingly, in one case (a spillover in Arizona), while we did not identify any previously characterised mutations in PB2 we did find PA M86I, which is at the same site as previously observed adaptive mutations M86V and T86I [26, 27]. The presence of uncharacterised mutations in the polymerase suggests (though does not prove) that novel mutations could be promoting mammalian replication in this outbreak. Although we did find potential mammalian adaptation mutations in the polymerase genes, we also confirmed with the Flu Mutation Explorer that none of the incursions into cattle showed any mammalian adaptations in HA.

These analyses demonstrate the way in which the Flu Mutation Explorer reduced the complex project of searching for signatures of mammalian adaptation across multiple viral genes during a rapidly developing outbreak to a simple task that could be performed robustly by users without specific bioinformatic training. They also demonstrate use of this tool to provide simple and specific data suitable for inclusion in influenza virus risk assessments.

## Discussion

Here we present the Flu Mutation Explorer, an interactive resource that integrates phylogenetic analyses with a curated catalogue of reported mammalian adaptation mutations to facilitate interpretation of IAV genomic variation. By placing mutations within their phylogenetic and host context, this resource enables rapid assessment of potential adaptive changes and complements existing influenza sequence analysis and surveillance platforms.

A central value of this resource lies in its manual curation of adaptive mutations, which prioritises traceability and interpretability. Each mutation is explicitly linked to primary literature, allowing users to evaluate the strength and context of the underlying evidence. This is particularly important in the field of influenza research, where the functional impact of mutations is often context-dependent, influenced by viral genetic background, host species, and experimental system. While automated approaches to mutation extraction are rapidly advancing, expert curation remains essential for resolving ambiguities and ensuring biological relevance. In this respect, the curated dataset presented here may also serve as a valuable benchmark for developing and validating machine learning approaches aimed at literature mining and genotype–phenotype mapping.

The integration of mutation data with phylogenetics provides important advantages for risk assessment and surveillance and our clustering approach allows for easy examination of IAV diversity. Rather than considering mutations in isolation, users can evaluate whether a mutation arises independently in multiple lineages, is associated with specific host transitions, or is restricted to particular clades. This is particularly relevant in the context of emerging zoonotic threats, such as recent H5N1 spillovers into cattle, where rapid identification and contextualisation of candidate adaptive mutations is essential. In such scenarios, Flu Mutation Explorer enables near real-time comparison of newly observed mutations against a curated knowledge base, supporting more informed interpretation of their potential significance.

Open sharing of influenza sequence data and associated metadata is essential for effective global surveillance and rapid assessment of emerging strains. However, the ability to analyse and interpret these data is often unevenly distributed, with many laboratories lacking access to specialised bioinformatics expertise or bespoke analytical pipelines. By providing an accessible and openly available resource that integrates curated mutation knowledge with phylogenetic context, Flu Mutation Explorer helps lower the barriers to analysis and supports broader participation in influenza risk assessment. This is particularly important for ensuring that countries contributing sequencing data can also rapidly evaluate the emergence potential of newly identified strains and place their findings within the context of global influenza diversity.

Despite these strengths, some limitations in the current version of this resource should be acknowledged. Clustering similar sequences means that mutations appearing only in an individual influenza sequence may in some cases not be visualised in a tree. Our mutation dataset is inherently influenced by biases in reported phenotypic data in the published literature, with a disproportionate focus on certain segments (notably PB2 and HA) and subtypes of public health concern (e.g. human seasonal H3N2). Mutations may only be significant in a specific sequence context as their influence can be impacted by the amino acid residues present at other sites, for example due to epistatic interactions. In particular, DMS screening for adaptive mutations, while generated at a large scale across a broad range of variants, is still very specific to the genotypes of the strains in which it was performed. Finally, while the application simplifies access to complex datasets, interpretation of the significance of any mutations identified in a strain still requires careful consideration of biological context and should not be viewed as definitive evidence of phenotype.

In summary, Flu Mutation Explorer provides a flexible and extensible platform for exploring influenza mutation data within an evolutionary framework. By bridging curated knowledge, genomic data, and phylogenetic context, it supports both routine surveillance and hypothesis-driven research. As IAVs continue to evolve and cross species barriers, integrative tools of this sort will be increasingly important for interpreting genetic variation and assessing its implications for public and animal health.

## Acknowledgements

We thank Scott Arkinson and Gary Cheung for IT support.

We acknowledge funding from the Medical Research Council (MRC) and Department for Environment, Food and Rural Affairs (Defra, UK) to the FluTrailMap-One Health Consortium [MR/Y03368X/1], from the MRC to the MRC-University of Glasgow Centre for Virus Research [MC_UU_00034/1, MC_UU_00034/5 and MC_UU_00034/6] and the University of Glasgow [MR/X502807/1], and from the Arts and Humanities Research Council to the University of Glasgow [AH/X003396/1]. T.P.P. is funded by funded by the BBSRC via the Pirbright Institute’s Strategic Programme Grants (ISPGs) [BBS/E/PI/230002A; BBS/E/PI/230002B].

## Author Contributions

LM: Data curation; Formal analysis; Investigation; Software; Methodology; Visualisation; Writing-Original draft preparation, reviewing and editing

DW: Software, Methodology

RG: Conceptualisation; Data curation

TP: Investigation

DLR: Conceptualisation; Supervision; Reviewing and Editing; Funding acquisition

JH: Conceptualisation, Methodology, Software, Visualisation, Supervision, Validation Writing-Original draft preparation, reviewing and editing

DHG: Investigation; Writing-Original draft preparation, reviewing and editing; Funding acquisition

EH: Conceptualisation, Supervision, Writing-Original draft preparation, reviewing and editing; Funding acquisition

## Supplementary Information

**Supplementary Figure 1:**
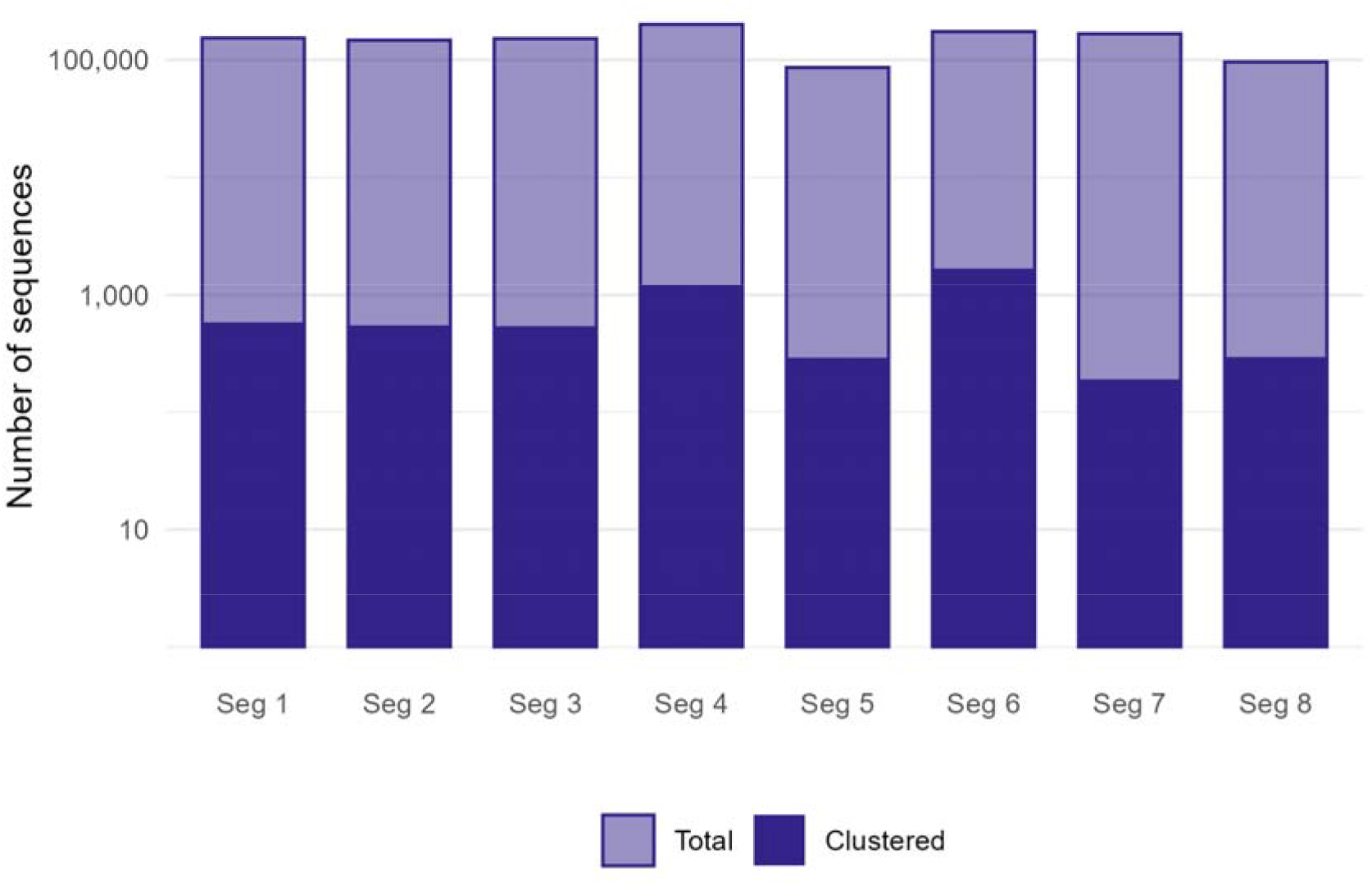
Overview of the IAV sequence database. Comparison of curated and clustered sequence counts for each IAV genome segment, as of June 8, 2026. Note that the y-axis is a logarithmic scale.

## Data Availability

The reference IAV sequences and alignments used in this study, and all scripts used for database pre-processing and code for the application, are available at https://github.com/centre-for-virus-research/Flu-Mutation-Explorer.

## References

1. Webster RG, Bean WJ, Gorman OT, Chambers TM, Kawaoka Y. Evolution and ecology of influenza A viruses. Microbiol Rev 1992;56:152–179.

2. Petrova VN, Russell CA. The evolution of seasonal influenza viruses. Nat Rev Microbiol 2018;16:47–60.

3. Long JS, Mistry B, Haslam SM, Barclay WS. Host and viral determinants of influenza A virus species specificity. Nat Rev Microbiol 2019;17:67–81.

4. WHO Tool for Influenza Pandemic Risk Assessment (TIPRA). https://www.who.int/publications/i/item/tool-for-influenza-pandemic-risk-assessment-(tipra)-2nd-edition (2020, accessed 17 June 2026).

5. Influenza Risk Assessment Tool (IRAT) Virus Descriptions and Report Summaries. https://www.cdc.gov/pandemic-flu/php/monitoring/virus-description.html (accessed 17 July 2026).

6. Technical risk assessment for avian influenza (human health): influenza A H5N1 2.3.4.4b. https://www.gov.uk/government/publications/avian-influenza-influenza-a-h5n1-risk-to-human-health/technical-risk-assessment-for-avian-influenza-human-health-influenza-a-h5n1-2344b (accessed 17 July 2026).

7. Global Influenza Surveillance and Response System (GISRS). https://www.who.int/initiatives/global-influenza-surveillance-and-response-system (accessed 17 July 2026).

8. Maurer-Stroh S. FluSurver: Real-time surveillance of influenza mutations. https://flusurver.bii.a-star.edu.sg (accessed 17 July 2026).

9. FluPhenotype: One-stop platform for early warnings of Influenza A viruses. https://flusurver.bii.a-star.edu.sg (accessed 17 July 2026).

10. Giussani E, Sartori A, Salomoni A, Cavicchio L, de Battisti C, et al. FluMut: a tool for mutation surveillance in highly pathogenic H5N1 genomes. Virus Evol 2025;11:veaf011.

11. Aksamentov I, Roemer C, Hodcroft E, Neher R. Nextclade: clade assignment, mutation calling and quality control for viral genomes. J Open Source Softw 2021;6:3773.

12. Kans J. Entrez® Direct: E-utilities on the Unix Command Line. https://www.ncbi.nlm.nih.gov/books/NBK179288/ (2013).

13. Altschul SF, Gish W, Miller W, Myers EW, Lipman DJ. Basic local alignment search tool. J Mol Biol 1990;215:403–410.

14. Steinegger M, Söding J. MMseqs2 enables sensitive protein sequence searching for the analysis of massive data sets. Nat Biotechnol 2017;35:1026– 1028.

15. Kalyaanamoorthy S, Minh BQ, Wong TKF, von Haeseler A, Jermiin LS. ModelFinder: fast model selection for accurate phylogenetic estimates. Nat Methods 2017;14:587–589.

16. Minh BQ, Schmidt HA, Chernomor O, Schrempf D, Woodhams MD, et al. IQ-TREE 2: New Models and Efficient Methods for Phylogenetic Inference in the Genomic Era. Mol Biol Evol 2020;37:1530–1534.

17. Suttie A, Deng Y-M, Greenhill AR, Dussart P, Horwood PF, et al. Inventory of molecular markers affecting biological characteristics of avian influenza A viruses. Virus Genes 2019;55:739–768.

18. Dadonaite B, Ahn JJ, Ort JT, Yu J, Furey C, et al. Deep mutational scanning of H5 hemagglutinin to inform influenza virus surveillance. PLOS Biol 2024;22:e3002916.

19. Soh YS, Moncla LH, Eguia R, Bedford T, Bloom JD. Comprehensive mapping of adaptation of the avian influenza polymerase protein PB2 to humans. eLife 2019;8:e45079.

20. Chang, W, Cheng, J, Allaire, J, Sievert, C, Schloerke, B, et al. shiny: Web Application Framework for R. https://github.com/rstudio/shiny (2026).

21. Sanderson T. Taxonium, a web-based tool for exploring large phylogenetic trees. eLife 2022;11:e82392.

22. Capelastegui F, Goldhill DH. H5N1 2.3.4.4b: a review of mammalian adaptations and risk of pandemic emergence. J Gen Virol 2025;106:002109.

23. Hutchinson EC, Denham EM, Thomas B, Trudgian DC, Hester SS, et al. Mapping the phosphoproteome of influenza A and B viruses by mass spectrometry. PLoS Pathog 2012;8:e1002993.

24. Burke DF, Smith DJ. A Recommended Numbering Scheme for Influenza A HA Subtypes. PLoS ONE 2014;9:e112302.

25. Burrough ER, Magstadt DR, Petersen B, Timmermans SJ, Gauger PC, et al. Highly Pathogenic Avian Influenza A(H5N1) Clade 2.3.4.4b Virus Infection in Domestic Dairy Cattle and Cats, United States, 2024. Emerg Infect Dis 2024;30:1335–1343.

26. Choi W-S, Baek YH, Kwon JJ, Jeong JH, Park S-J, et al. Rapid acquisition of polymorphic virulence markers during adaptation of highly pathogenic avian influenza H5N8 virus in the mouse. Sci Rep 2017;7:40667.

27. Welkers MRA, Pawestri HA, Fonville JM, Sampurno OD, Pater M, et al. Genetic diversity and host adaptation of avian H5N1 influenza viruses during human infection. Emerg Microbes Infect 2019;8:262–271.

